# Too much sugar makes plants ‘pregnant’: maternal sucrose signals fertilization in Arabidopsis seeds

**DOI:** 10.1101/2025.05.16.654273

**Authors:** Wenjia Xu, Miryam Iannaccone, Dennys-Marcela Gomez-Paez, Sandrine Choinard, Jing Lu, Rozenn Le Hir, Sylvie Dinant, Lothar Kalmbach, Regina Feil, John E. Lunn, Christian Meyer, Enrico Magnani

## Abstract

Reproduction is deemed successful if it ensures the perpetuation of the species. Nonetheless, evolution has optimized reproductive efficiency across a diverse range of strategies. Animal vivipary, the process in which embryos develop inside the parent’s body, allows for larger offspring and better protection. This reproductive strategy relies on a fertilization signal sent to the maternal tissues to establish an embryo-supportive environment, a process known as “maternal recognition of pregnancy”. Despite being a strictly animal-only term, plants independently evolved a conceptually similar strategy: the seed habit, wherein the embryo is nourished and protected by surrounding maternal tissues. We discovered that sucrose is sufficient to trigger a fertilization-independent response in the maternal tissues of Arabidopsis ovules. Upon fertilization, the release of Polycomb-mediated repression promotes the symplastic and apoplastic flow of sucrose into the seed by modulating auxin and trehalose 6-phosphate signaling. The sucrose signal is then perceived by the TARGET OF RAPAMYCIN kinase and transduced into a gibberellin response to initiate the differentiation of the seed maternal tissues. This work revealed a zygotic-maternal interplay of hormonal and sugar signaling that drives the plant “maternal recognition of fertilization”, incorporating energetic cues into developmental pathways.

## Introduction

Fertilization is the fusion of male and female gametes to form a zygote, the next sporophytic generation. This fundamental process is conserved across all species that reproduce sexually. However, a yeast zygote is released in the environment while a human embryo requires the successful implantation in the surrounding maternal tissues to achieve its proper development. Fertilization in viviparous animals, where the embryo grows inside the parent, evolved signaling functions to promote the development of a maternal milieu supportive of embryogenesis, a process referred to as “maternal recognition of pregnancy” (*1, 2*). The evolution of zygote-maternal communication allowed higher survival rate, better adaptation to different environments, greater diversification and paved the way to increased parental investment in offspring, influencing social structures and mating dynamics within populations. In mammals, maternal recognition of pregnancy begins with a signal sent by the embryo to the maternal corpus luteum to preserve its function and structure, which would otherwise regress at the end of the estrous or menstrual cycle (*3*). For example, the primate conceptus produces chorionic gonadotropin, a signal that maintains progesterone production in the corpus luteum, which in turn supports the endometrium during early embryonic development, implantation, placentation, and the growth of the fetoplacental unit.

In the 17th century, botanists like Nehemiah Grew and Rudolf Jakob Camerarius, made significant contributions to the understanding of plant reproduction, comparing plant reproductive structures to those of animals (*4, 5*). This concept was intuitively captured by artists such as the American surrealist painter Helen Lundeberg, who drew a striking parallel between the human uterus and a cherry in her 1934 artwork *Plant and Animal Analogies* (UCI Museum and Institute for California Art, USA). The reproductive strategy of seed plants is indeed conceptually similar to that of viviparous animals, as the plant embryo develops inside maternal tissues that protect and nourish it. As a consequence, seeds independently evolved a signaling pathway equivalent to the animal maternal recognition of pregnancy that informs maternal tissues when fertilization occurs. This process has been partially elucidated in Arabidopsis, where a double fertilization event generates the embryo, the next sporophytic generation, and the endosperm, which supports the embryo’s growth (*6*). The endosperm, through the action of the AGAMOUS LIKE 62 (AGL62) MADS domain transcription factor, induces a hormonal cascade of auxin and gibberellin that promotes the development of the surrounding maternal tissues (*7*). *agl62* mutant seeds are unable to transduce the fertilization signal, resulting in undeveloped maternal tissues (*8*). By contrast, unfertilized ovules treated with auxin or gibberellin differentiate prematurely (*7*). Two phenotypic markers of maternal response are the expansion of the seed coat, which surrounds the fertilization products, and the synthesis of proanthocyanidins (PAs) in the endothelium, the innermost layer of the seed coat (Fig. 1 A and B). Before fertilization, members of the FERTILIZATION INDEPENDENT SEED (FIS)-Polycomb Group (PcG) proteins repress the development of seed maternal tissues (*8*). Mutants for the *FERTILIZATION-INDEPENDENT ENDOSPERM* (*FIE*) or *MULTICOPY SUPPRESSOR OF IRA1* (*MSI1*) *FIS* genes produce enlarged ovules containing PAs in the absence of fertilization (*8, 9*).

**Figure 1.**
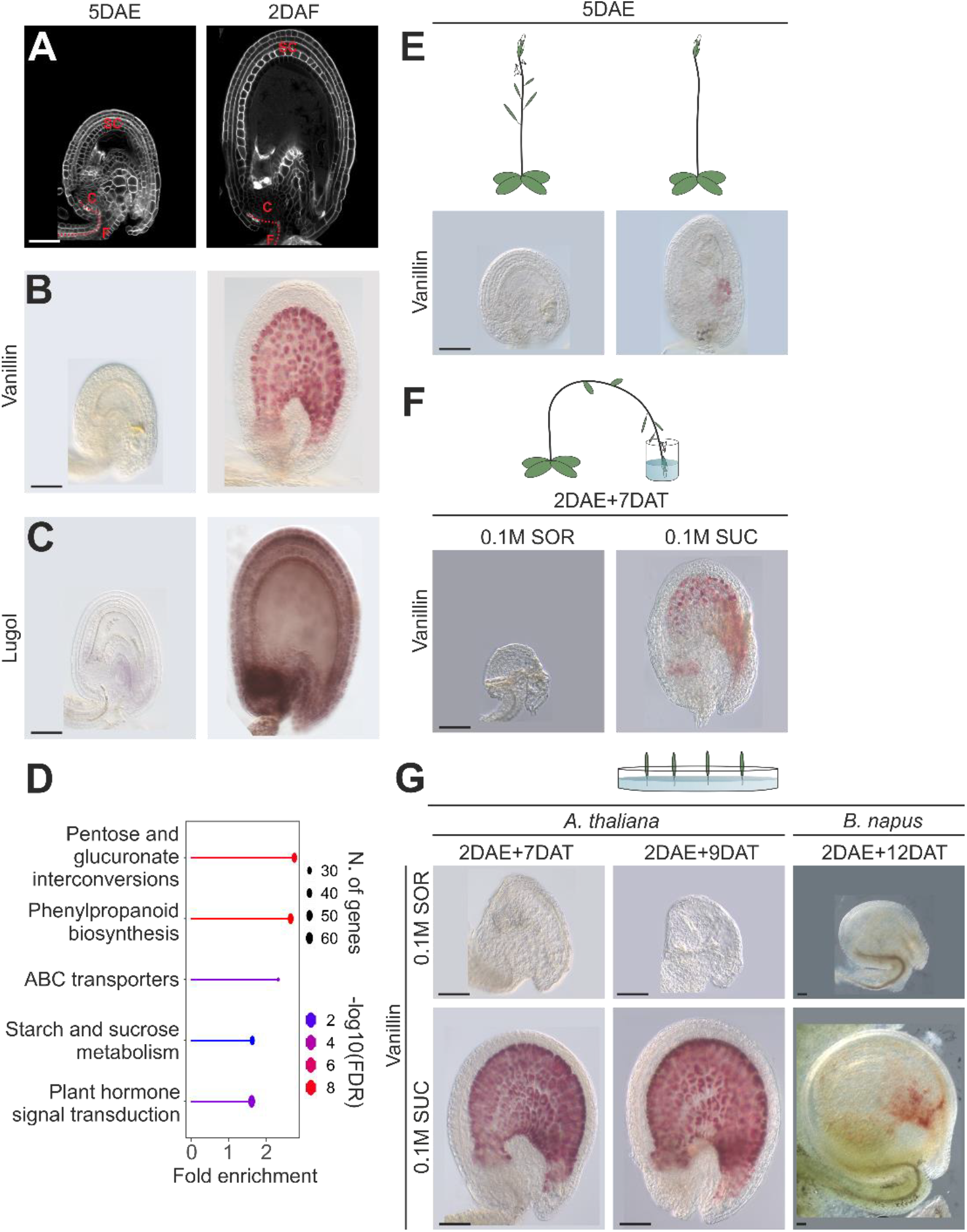
Sucrose induces a fertilization response in maternal tissues. **(A)** Confocal images of an ovule and seed stained with SR2200. Dotted red lines highlight the vascular tissues. F, funiculus; C, chalaza; SC, seed coat. **(B-C)** Differential interference contrast (DIC) microscopy images of ovules and seeds stained with vanillin (to detect PAs, **B**) or Lugol’s solution (to detect starch, **C**). **(D)** Enriched GO terms for genes differentially expressed in seeds across fertilization (log2 fold change threshold +/-1). **(E)** DIC images of ovules obtained from untrimmed or trimmed inflorescences and stained with vanillin. **(F)** DIC images of ovules obtained from inflorescences dipped in a sorbitol or sucrose solution and stained with vanillin. **(G)** DIC images of ovules obtained from pistils grown *in vitro* in a sorbitol or sucrose medium and stained with vanillin. DAE, days after emasculation; DAF, days after flowering; DAT, days after treatment; SOR, sorbitol; SUC, sucrose; Bars = 50 µm.

Zygotic-maternal communication must also guarantee the balanced allocation of resources to the progeny, which is fundamental to the success of reproduction (*10*). Maternal tissues must correctly sense fertilization to provision the offspring with enough nutrients to ensure its vigor while not over-depleting the parent and, thereby, endanger its survival. This is particularly true in sessile organisms, such as plants, which are more influenced by environmental conditions and nutrient availability than motile organisms. Building on this premise, we hypothesized the existence of a maternal checkpoint within the fertilization signaling pathway, integrating information about the plant’s energetic status. Sucrose, ultimately derived from photosynthesis, is the main source of carbon and energy for growing sink organs, such as seeds. We discovered that maternal sucrose is sufficient to trigger a fertilization signal in Arabidopsis seeds. Our data unveil a complex signaling pathway of hormonal and sugar cues between zygotic and maternal tissues, responsible for the plant “maternal recognition of fertilization”.

## Results

### Maternal sucrose is sufficient to induce a fertilization signal

In Arabidopsis, seed coat expansion and PA biosynthesis are hallmarks of maternal response to fertilization (Fig. 1A and B). We observed that these developmental changes are closely linked to the accumulation of starch, the primary form of storage-carbohydrate in plants (Fig. 1C). Consistently, we obtained significant enrichments in GO annotations for ‘starch and sucrose metabolism’ together with ‘phenylpropanoid biosynthesis’ (PAs are part of the phenylpropanoids pathway) in transcriptomic analyses conducted in seeds across fertilization (Fig. 1D and Supplementary Table 1). In line with these results, seeds displayed higher levels of soluble sugars (sucrose, glucose, and fructose) than unfertilized ovules (Supplementary Fig. 1), indicating that fertilization stimulates the uptake of sugars. By contrast, we detected less starch in *agl62* mutant seeds, which fail to generate a fertilization signal and lack a maternal response (*7, 8*), when compared to the wild type (Supplementary Fig. 2).

Sucrose, the major form of transport-sugar in plants, is not only a source of energy and carbon but also a well-established signaling molecule (*11-14*). This dual role led us to hypothesize that sucrose might be involved in the fertilization signaling pathway beyond its metabolic function. To test this hypothesis, we trimmed developing inflorescences off their siliques to direct more assimilates to the reproductive organs. As a result, we observed bigger ovules exhibiting PA deposition in the absence of fertilization, when compared to untrimmed inflorescences (Fig. 1E and Supplementary Fig. 3). In addition, we supplemented unfertilized ovules with 0.1 M sucrose, which is well within the physiological concentration of sucrose delivered in the phloem (*15, 16*), in *in planta* and *in vitro* experiments (Fig. 1F and G, see Materials and Methods). In both experimental settings, sucrose stimulated PA deposition and growth of the seed coat similar to what observed in seeds (Fig. 1F and G and Supplementary Fig. 3 and 4), while glucose, fructose, or a combination of the two produced a milder response (Supplementary Fig. 3). Conversely, equal molarity of sorbitol, a non-metabolizable sugar, did not elicit developmental effects (Fig. 1G and Supplementary Fig. 3 and 4). We obtained similar results with *B. napus* ovules, thus indicating that this sucrose response is conserved among Brassicaceae (Fig. 1G). Altogether, these data indicate that sucrose is sufficient to trigger a fertilization response in ovule maternal tissues.

### Fertilization promotes the symplastic and apoplastic movement of sugars into the seed

The finding that sucrose triggers a fertilization-independent response in ovules highlights the importance of tightly regulating its flow. Sucrose is transported from the photosynthetic tissues of the plant to the seed via the phloem. The vascular tissues extend through the funiculus and terminate in the chalazal region of the seed, the nutrient unloading zone (Fig. 1A). Further transport occurs either symplastically, via plasmodesmata, or apoplastically, through plasma membrane transporters. (*17, 18*). To investigate whether fertilization regulates the symplastic connections in the seed, we tracked the movement of carboxyfluorescein (CF), a fluorescent tracer for symplastic transport (*19*). Three hours after treatment, CF fluorescence spread throughout the chalaza and seed coat of young seeds, whereas it remained confined to the phloem in unfertilized ovules (Fig. 2A). This result indicates that fertilization facilitates phloem unloading in the chalaza. Since symplastic movement is hindered by callose deposition at sieve pores and plasmodesmata (*20*), we analyzed the expression of *CALLOSE SYNTHASE* (*CALS*) genes in the ovule through *in silico* (*21*) and promoter-reporter gene analyses. We identified the paralogue genes *CALS6* and *7* (*22-25*) as being expressed in the funiculus and chalaza (Fig. 2B and Supplementary Fig. 5). Double *cals6;7* mutant ovules accumulated more soluble sugars and starch than the wild type and, as predicted, developed larger seed coats rich in PAs (Fig. 2C and Supplementary Fig. 1 and 3). By contrast, the induction of *CALS3* expression (*26*) in unfertilized ovules cultured on sucrose inhibited starch accumulation and the development of the maternal tissues (Fig. 2D and Supplementary Fig. 3). In line with this model, our transcriptomic analyses revealed that fertilization promotes the expression of several *GLUCANASE* genes (Fig. 2E and Supplementary Table 1), four of which have known callose hydrolases activity, that facilitate sugar uptake as recently shown by Liu and coworkers (*27*).

**Figure 2.**
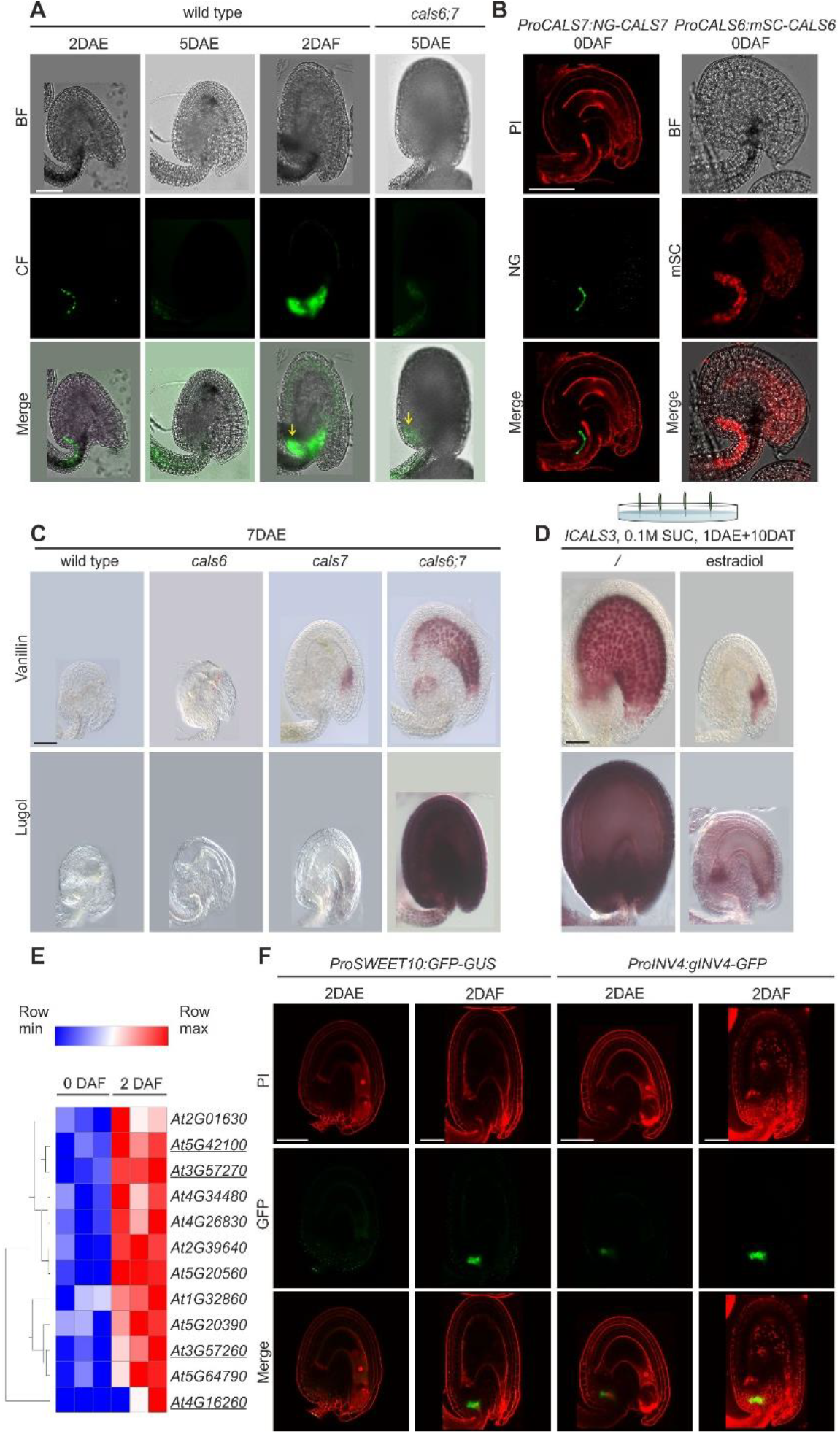
Fertilization promotes sugar symplastic and apoplastic transport. **(A)**Confocal images of wild type and mutant ovules and seeds obtained from pistils and siliques loaded with carboxyfluorescein diacetate. Yellow arrows indicate carboxyfluorescein (CF) diffusion in the chalaza. **(B)** Confocal images of ovules from *CALS6* or *CALS7* promoter-reporter lines. **(C)** DIC images of wild type and mutant ovules stained with vanillin or Lugol’s solution. **(D)** DIC images of *ICALS3* mock and induced ovules stained with vanillin or Lugol’s solution. **(E)** Heatmaps (three replicates per time point) showing the expression of *GLUCANASE* genes up-regulated after fertilization. The underlined genes have known callose hydrolase activity. **(F)** Confocal images of ovules and seeds from *SWEET10* and *INV4* promoter-reporter lines. /, mock treatment; DAE, days after emasculation; DAF, days after flowering; DAT, days after treatment; BF, bright field; mSC, mScarlet; NG, Neongreen; PI, propidium iodide; SUC, sucrose; Bars = 50 µm.

Apoplastic sugar transport has been shown to play a crucial role in both early and late stages of seed development (*18, 28*). Sucrose moves to the apoplast through SWEET facilitators (*29*) following a concentration gradient, which is maintained by cell wall invertases that convert sucrose into hexoses (*30*), or by active uptake by SUT/SUC type sucrose:H^+^ symporters (*31-33*). We identified the sucrose facilitator *SWEET10* (*34*) and cell wall *INVERTASE4* (*35*), as being upregulated by fertilization in the chalazal region of the seed (Fig. 2F and Supplementary Fig. 5 and 6). Altogether, these results indicate that fertilization transforms the seed into a sink for sucrose, which in turn signals fertilization to the maternal tissues.

### Fertilization promotes sugar transport by modulating T6P and auxin signaling

Our metabolic analyses showed that fertilization induces the accumulation of trehalose 6-phosphate (T6P) (Fig. 3A), an intermediate in the synthesis of the sugar trehalose. T6P closely mirrors sucrose levels (*36*) and functions as a signaling molecule in diverse developmental pathways (*36-39*). Consistently, we detected significantly less T6P in *agl62* seeds, when compared to the wild type (Fig. 3B). We profiled the expression of trehalose metabolic genes, *TREHALOSE-6-PHOSPHATE SYNTHASE*S (TPSs) and *PHOSPHATASES* (TPPs), and identified a transcriptional regulation in response to fertilization (Fig. 3C and Supplementary Table 1). Among them, *TPS1*, a well characterized T6P biosynthetic gene (*37, 38, 40*), was expressed in the funiculus and chalaza regions (Fig. 3D and Supplementary Fig. 5). To study the role of T6P in the fertilization signaling process, we rescued the embryo lethal *tps1/tps1* mutant using a dexamethasone-inducible *TPS1* expression system (*ITPS;tps1*) (*37*) and stopped induction after flowering. We observed that 18% of *ITPS;tps1* seeds failed to grow and to produce PAs all along the seed coat (Fig. 3E and Supplementary Fig. 3), similar to what detected in the *agl62* mutant (Supplementary Fig. 2) (*8*). These underdeveloped seeds did not accumulate starch (Fig. 3E) and were impaired in CF movement to the chalaza (Fig. 3F), indicating a direct role for T6P in facilitating sugar transport.

**Figure 3.**
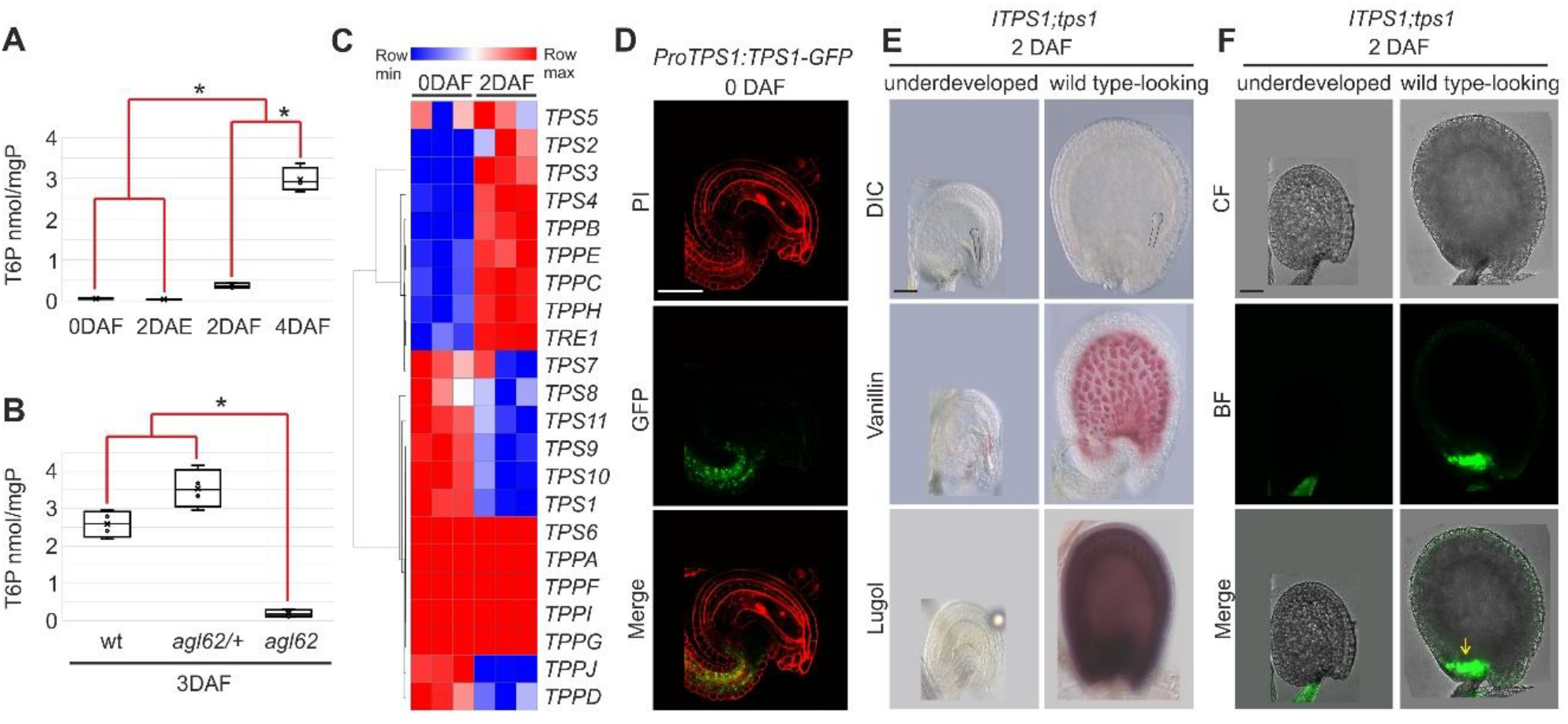
T6P promotes seed symplastic transport. **(A-B)** Quantification of T6P levels in wild type and *agl62* ovules and seeds. In the box and whisker chart, the rectangles represent the interquartile ranges (median excluded), the mid-lines represent the medians and the whiskers represent the minimum and maximum values within 1.5 times the interquartile range. Asterisks indicate statistically significant difference (Student’s t-test, P<0.05, n=4 biological replicates). **(C)** Heatmaps (three replicates per time point) showing the expression of trehalose biosynthetic and related genes across fertilization. **(D)** Confocal images of ovules from the *TPS1* promoter-reporter line. **(E)** DIC images of underdeveloped or wild type-looking *ITPS1;tps1* seeds stained with vanillin or Lugol’s solution. **(F)** Confocal images of underdeveloped or wild type-looking *ITPS1;tps1* seeds obtained from siliques loaded with carboxyfluorescein diacetate. The yellow arrow indicates carboxyfluorescein (CF) diffusion in the chalaza. mgP, mg of protein; DAE, days after emasculation; DAF, days after flowering; BF, bright field; PI, propidium iodide; T6P, trehalose 6-phosphate; Bars = 50 µm.

Fertilization has also been shown to trigger an auxin response in maternal tissues (*7*). As previously shown, treatments with auxin stimulated cell expansion and PA deposition in the integuments of unfertilized ovules (Fig. 4A) (*7*). In addition, auxin induced the accumulation of starch (Fig. 4A) and enhanced the maternal response to sucrose *in vitro* (Fig. 4B and Supplementary Fig. 3), suggesting its involvement in regulating sugar transport. Consistent with this hypothesis, auxin upregulated the expression of *SWEET10* and *INV4*, as detected by promoter-GFP lines (Fig. 4C and Supplementary Fig. 6), and led to increased levels of sucrose, glucose and fructose (Fig. 4D). Furthermore, it promoted the synthesis of T6P in unfertilized ovules (Fig. 4E). Nevertheless, *in vitro* experiments revealed that auxin elevated T6P levels only in the presence of sucrose, while no such effect was observed in a sorbitol medium (Fig. 4E). This suggests that auxin indirectly regulates T6P synthesis by enhancing sucrose availability. Conversely, sucrose did not stimulate an auxin response in unfertilized ovules carrying a *DR5v2:GFP* reporter system (*41*) (Fig. 4F) and did not trigger division of the central cell (Fig. 4F), a phenotype induced by auxin treatments (*42*). These results indicate that auxin facilitates the apoplastic uptake of sucrose into the seed and indirectly T6P synthesis.

**Figure 4.**
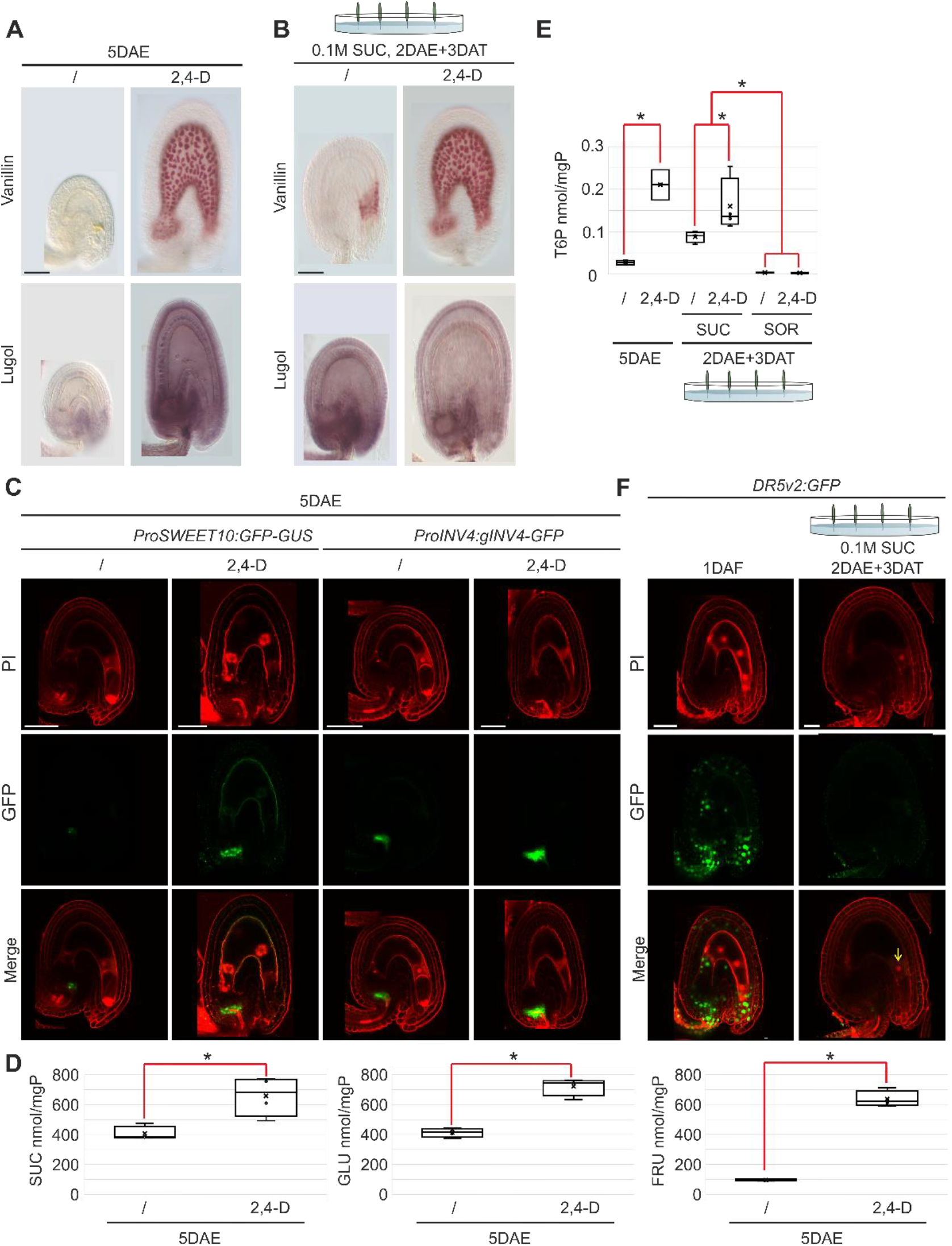
Auxin promotes seed apoplastic transport. **(A-B)** DIC images of unfertilized ovules treated with the synthetic 2,4-D auxin *in planta* **(A)** and *in vitro* **(B)**stained with vanillin or Lugol’s solution. **(C)** Confocal images of ovules from *SWEET10* or *INV4* promoter-reporter lines. **(D)** Quantification of sucrose, glucose, and fructose levels in ovules treated with auxin. In the box and whisker chart, the rectangles represent the interquartile ranges (median excluded), the mid-lines represent the medians and the whiskers represent the minimum and maximum values within 1.5 times the interquartile range. Asterisks indicate statistically significant difference (Student’s t-test, P<0.05, n=4 biological replicates). **(E)** Quantification of T6P levels in ovules treated with auxin. In the box and whisker chart, the rectangles represent the interquartile ranges (median excluded), the mid-lines represent the medians and the whiskers represent the minimum and maximum values within 1.5 times the interquartile range. Asterisks indicate statistically significant difference (Student’s t-test, P<0.05, n=4 biological replicates). **(F)** Confocal images of *DR5V2:GFP* ovules and seeds. The central cell is highlighted by a yellow arrow. /, mock treatment.; mgP, mg of protein; DAE, days after emasculation; DAF, days after flowering; DAT, days after treatment; FRU, fructose; GLU, glucose; PI, propidium iodide; SOR, sorbitol; SUC, sucrose; T6P, trehalose 6-phosphate; Bars = 50 µm.

### FIE represses T6P and auxin signaling

The FIS group of PcG proteins prevents the development of maternal tissues prior to fertilization (*43*). To test whether PcG repression influences sugar transport, we analyzed the role of FIE, which is expressed in all maternal tissues (Fig. 5A and Supplementary Fig. 5) (*9, 44*). The *fie* mutation is haploinsufficient and emasculated *fie/+* flowers produce both wild type-looking and enlarged ovules that accumulate PAs (Fig. 5B) (*8*). In line with our previous results, we detected more starch in *fie/+* enlarged ovules when compared to *fie/+* wild type-looking ovules (Fig. 5B). The *fie* mutation induced the production of T6P even when grown exclusively on sorbitol, indicating that FIE represses T6P synthesis in a sucrose-independent fashion (Fig. 5C). Furthermore, we detected auxin response in the funiculus and chalaza of enlarged *fie/+* ovules carrying a *DR5v2:GFP* reporter system, whereas no signal was visible in *fie/+* wild-type looking ovules (Fig. 5D). Overall, these results indicate that FIE represses T6P and auxin pathways, thus preventing the fertilization-independent flow of sucrose into the ovule.

**Figure 5.**
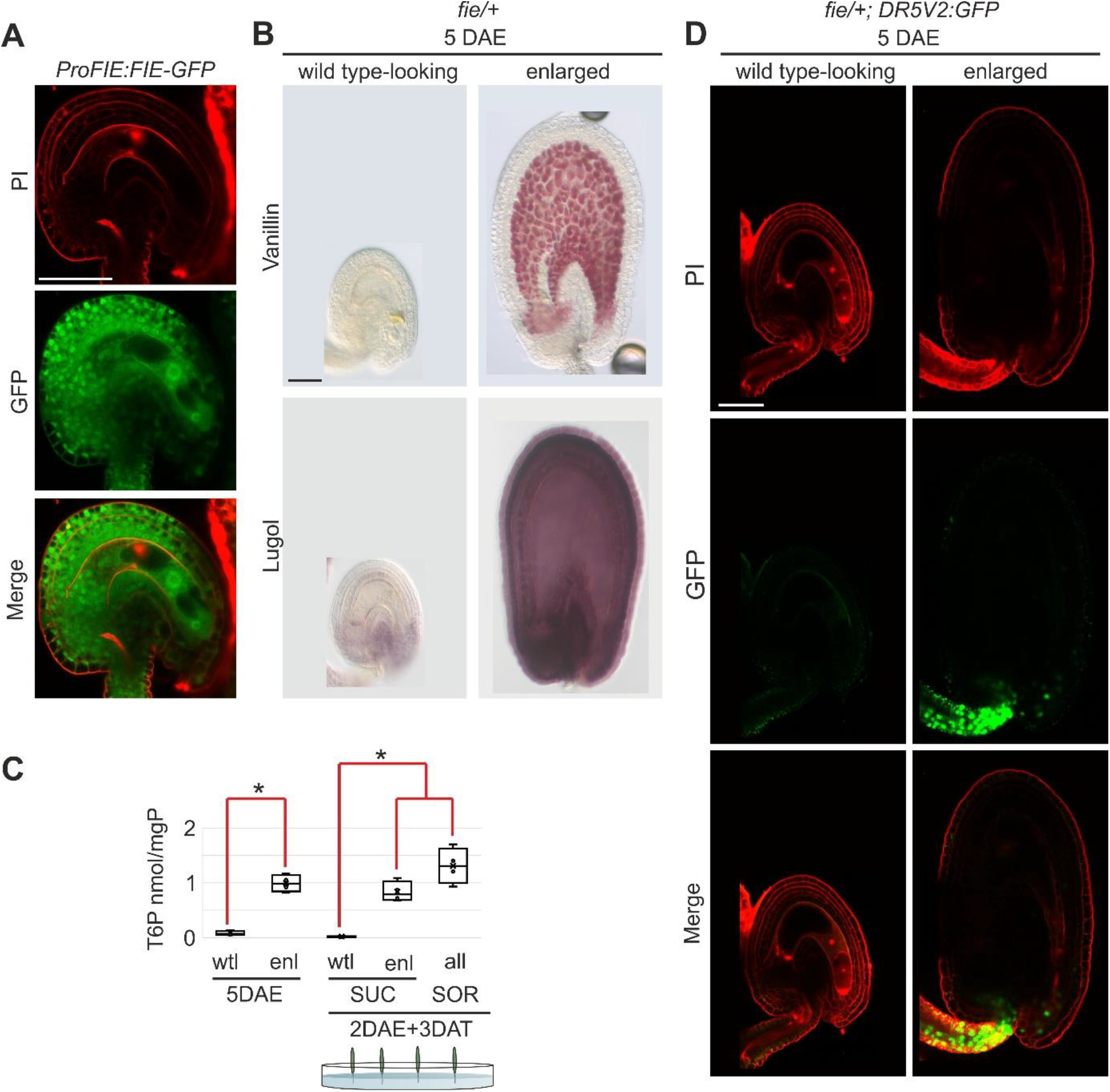
FIE represses T6P and auxin signaling before fertilization. **(A)** Confocal images of ovules from *FIE* promoter-reporter line. **(B)** DIC images of *fie/+* unfertilized ovules stained with vanillin or Lugol’s solution. **(C)** Quantification of T6P levels in *fie/+* wild type-looking (wtl) and enlarged (enl) unfertilized ovules. In the box and whisker chart, the rectangles represent the interquartile ranges (median excluded), the mid-lines represent the medians and the whiskers represent the minimum and maximum values within 1.5 times the interquartile range. Asterisks indicate statistically significant difference (Student’s t-test, P<0.05, n=4 biological replicates). **(D)** Confocal images of *fie/+*;*DR5V2:GFP* ovules. mgP, mg of protein; DAE, days after emasculation; DAT, days after treatment; PI, propidium iodide; SOR, sorbitol; SUC, sucrose; T6P, trehalose 6-phosphate; Bars = 50 µm.

### Sucrose triggers a gibberellin response

Gibberellins have been shown to act downstream of auxin in the fertilization signaling pathway (*7*). When gibberellic acid 3 (GA3) was applied to unfertilized ovules, it promoted PA deposition and growth of the seed coat (Fig. 6A), although the latter response was less pronounced compared to sucrose or auxin treatments (Fig. 1G and 4A). Nevertheless, GA3 did not induce the accumulation of starch (Fig. 6A), suggesting that gibberellins act downstream of sucrose. To test this hypothesis we used the DELLA reporter *ProRGA:GFP-RGA*, an inverse-fluorescent marker for active gibberellin signaling (*45*). Sucrose triggered a gibberellin response, observed as a decrease in GFP fluorescence, within the seed coat of unfertilized ovules cultured *in vitro* (Fig. 6B). We observed the strongest response in the endothelium, which synthetizes PAs, whereas the outermost integument remained unaffected (Fig. 6B and C). To further challenge our model, we repressed gibberellin signaling by over-expressing the *DELLA* genes *GAINSENSITIVE* (GAI, *Pro35S:GAI*) or its mutated version *GAI∆17* (*Pro35S:GAI∆17*), which is insensitive to gibberellin induced proteolysis (*46*). Both over-expressing lines inhibited PA deposition, but not starch accumulation, in fertilized seeds *in planta* (Fig. 6D and Supplementary Fig. 3), and unfertilized ovules treated with sucrose *in vitro* (Fig. 6E and Supplementary Fig. 3). Altogether, these data suggest that sucrose triggers a gibberellin response in the internal cell layers of the seed coat, leading to PA deposition.

**Figure 6.**
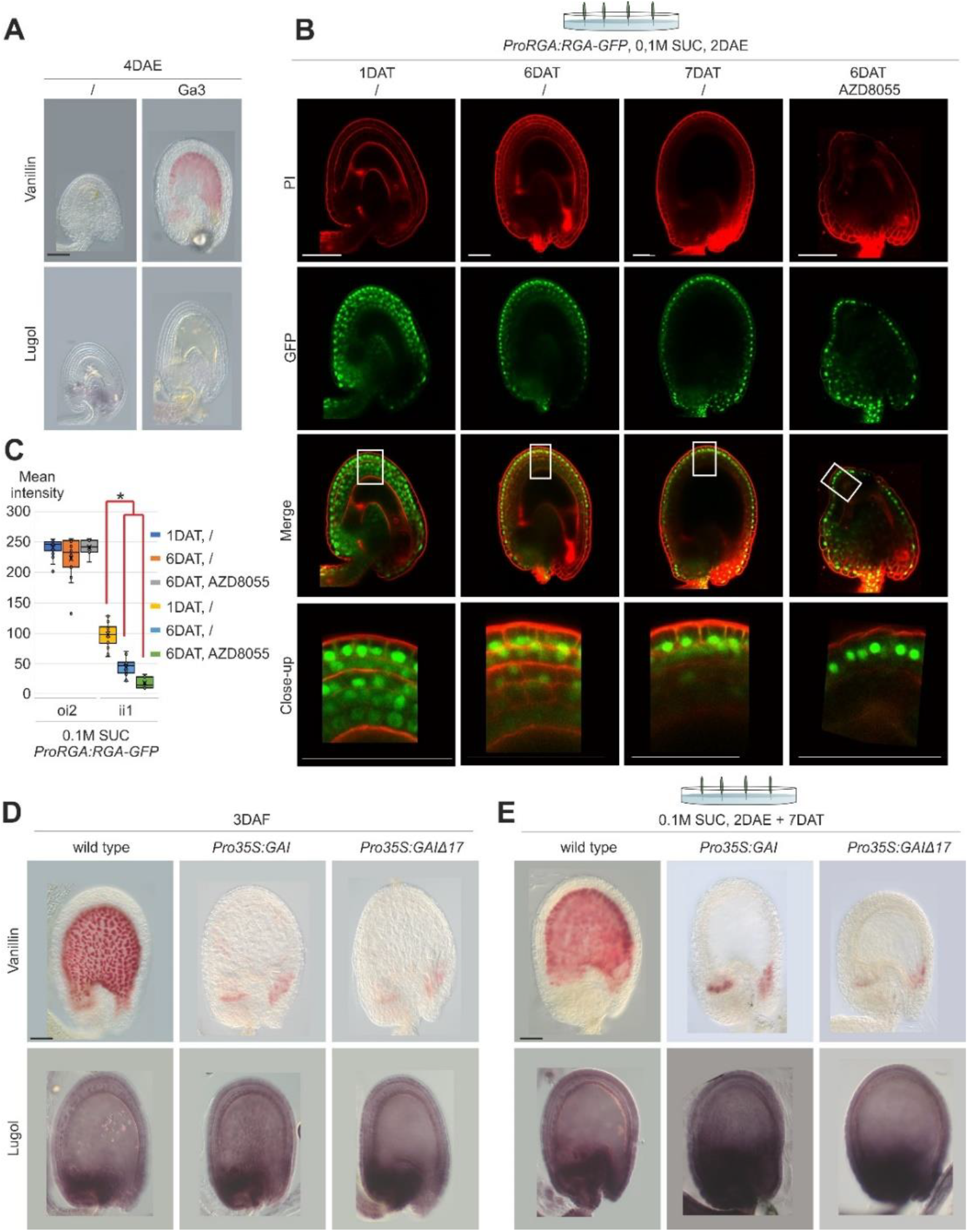
Sucrose induces a gibberellin response in seed maternal tissues. **(A)** DIC images of unfertilized ovules treated with gibberellic acid 3 (GA3) stained with vanillin or Lugol’s solution. **(B)** Confocal images of *proRGA:RGA-GFP* unfertilized ovules. **(C)** Quantification of GFP intensity in the outer integument 2 (oi2) and inner integument 1 (ii1) integuments of *proRGA:RGA-GFP* unfertilized ovules. In the box and whisker chart, the rectangles represent the interquartile ranges (median excluded), the mid-lines represent the medians and the whiskers represent the minimum and maximum values within 1.5 times the interquartile range. Asterisks indicate statistically significant difference (Student’s t-test, P<0.05, n>7). **(D-E)** DIC images of wild type, *Pro35S:GAI*, and *Pro35S:GAI∆17* seeds and ovules stained with vanillin or Lugol’s solution. /, mock treatment; DAE, days after emasculation; DAF, days after flowering; DAT, days after treatment; PI, propidium iodide; SUC, sucrose; Bars = 50 µm.

### TOR transmits the sucrose signal

Sugars are known to activate the TARGET OF RAPAMYCIN (TOR) kinase, a central regulator of energy availability that links external and internal cues to control growth in eukaryotes (*47, 48*). A T-DNA-mediated translational fusion of *TOR* with the *GUS* reporter gene (*47*) showed *TOR* expression in all maternal tissues of ovules and young seeds (Fig. 7A and Supplementary Fig. 5). To test TOR function in the fertilization signaling pathway, we analyzed an inducible *TOR* artificial RNAi (*IRNAi-TOR*) line (*49*), as the *tor* null mutation is embryo lethal (*47*). The induction of *IRNAi-TOR* inhibited PA deposition and growth of the seed coat after fertilization (Fig. 7B and Supplementary Fig. 3), when compared to an inducible *GUS* line (*IGUS*) as control. Likewise, TOR inhibitors, such as TORIN2 and AZD8055 (*50-52*), as well as mutations in the TOR interactor *REGULATORY-ASSOCIATED PROTEIN OF TOR 1B* (*RAPTOR*) gene (*53*), repressed the maternal response to sucrose of unfertilized ovules cultured *in vitro* (Fig. 7C and Supplementary Fig. 3). By contrast, inhibition of *TOR* expression or function did not significantly affect starch accumulation (Fig. 7B and C), suggesting that TOR does not affect sucrose transport. Finally, we tested whether TOR transduces the sucrose signal by modulating the gibberellin pathway. The treatment of unfertilized ovules with AZD8055 did not suppress the gibberellin response, as measured with the *ProRGA:GFP-RGA* reporter (Fig. 6B). Furthermore, GA3 treatments failed to restore a wild type maternal response in unfertilized ovules cultured *in vitro* with sucrose and AZD8055 (Fig. 7C and Supplementary Fig. 3). Altogether, these data suggest that TOR acts in concert with gibberellins to execute sugar cues in response to fertilization.

**Figure 7.**
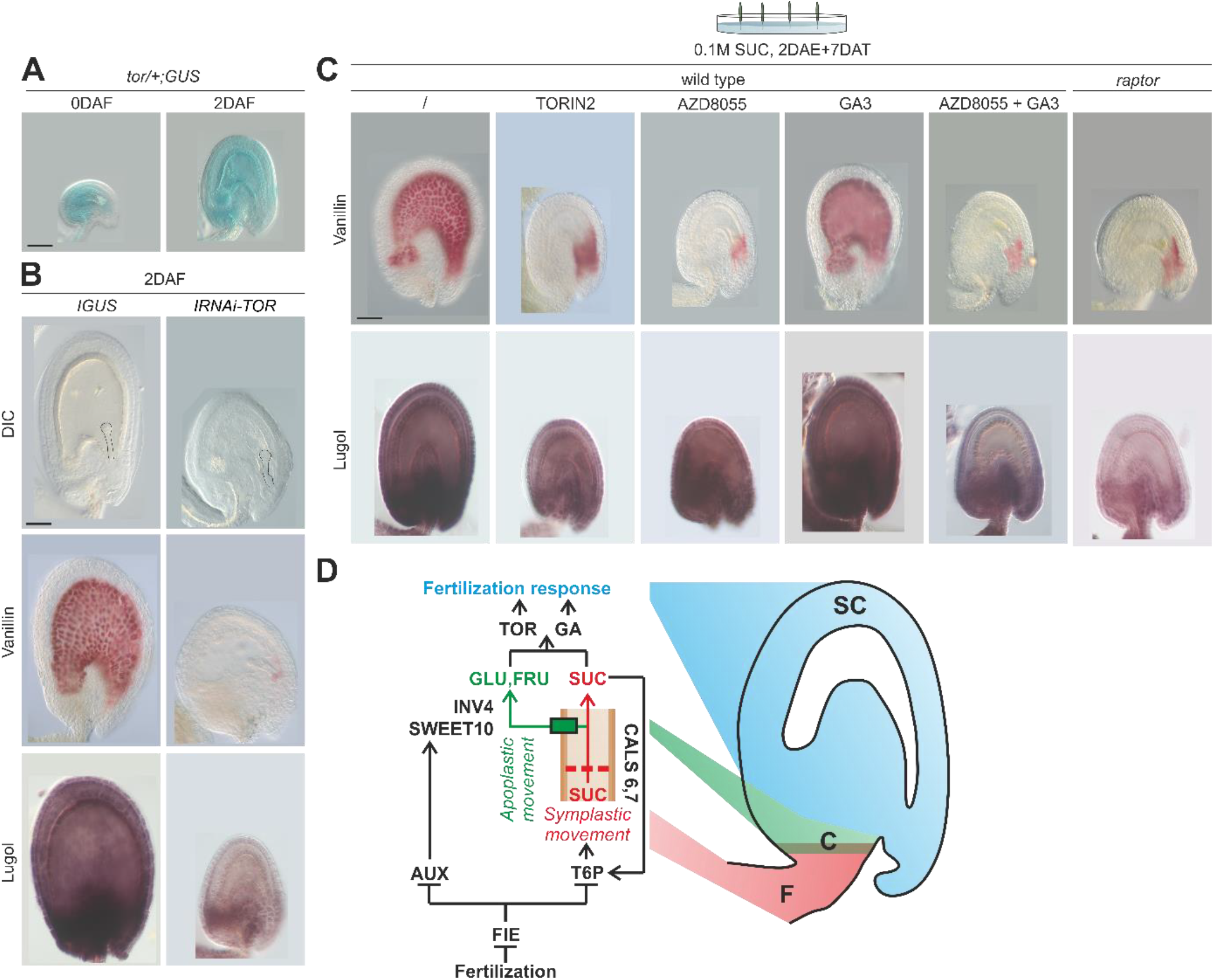
TOR transduces the sucrose signal in seed maternal tissues. **(A)** DIC images of a *tor/+;GUS* ovule and seed. **(B)** DIC images of *IRNAi-TOR* and *IGUS* control seeds stained with vanillin or Lugol’s solution. The embryo is outlined in black. **(C)** DIC images of unfertilized wild type or mutant ovules grown *in vitro* in the presence or not of TOR inhibitors. **(D)** Model for sucrose signaling during fertilization. /, mock treatment; DAE, days after emasculation; DAF, days after flowering; DAT, days after treatment; SUC, sucrose; GLU, glucose; FRU, fructose; F, funiculus (red); C, chalaza (green); SC, seed coat (cyan); Bars = 50 µm.

## Materials and methods

### Genetic Materials

Col-0, *cal6-1* (SALK_019534), *cal7-1* (SALK_048921), *cal6;7, agl62-2* (SALK_022148), *fie-12* (GK-362D08), *raptor78* (SALK_078159). *ICALS3* line was described in (*26*). *tor/+;GUS* line was described in (*47*). *IRNAi-TOR* and *IGUS* lines were described in (*49*).

*ITPS1;tps1* line was described in (*37*). ProCALS7:NG-CALS7 line was described in (*25*). *DR5V2:GFP* line was described in (*41*). *ProTPS1:TPS1-GFP* line was described in (*40*). The *ProFIE:FIE-GFP* line was described in (*44*). *ProRGA:RGA-GFP* line was described in (*45*). *Pro35SGAI* and *Pro35S:GAI∆17* lines were described in (*46*).

### Cloning

*ProCALS6:mSC-CALS6* was assembled through a 3x multisite Gateway reaction using the entry clones pENTR-L4-pCALS6-R1, pENTR-L1-mSC-L2 and pR2-CALS6-L3. To generate the pCALS6 entry clone the 2463 bp region immediately upstream of the ATG of *CALS6* (AT3G59100) was PCR amplified (primers 5’TCTCGTTCAGCTTTTTTGTACAAACTTGTatccaccgtgaatcgaaactataa3’ and 5’-ATGCCAACTTTGTATAGAAAAGTTGTTgtatgtggtattcctctgcaca-3’) with matching overlapping regions for Gibson Assembly with a PCR fragment containing an L4-R1 vector backbone (amplified with primer 5’AACAACTTTTCTATACAAAGTTGGCAT3’ and 5’ACAAGTTTGTACAAAAAAGCTGAACGAGA3’). To generate the genomic *CALS6* clone, 3 fragments of approximately 3.5 kb covering the entire genomic *CALS6* region were PCR amplified (using primers 5’TTCAGCTTTCTTGTACAAAGTGGTTATGGAAGCTAGTTCGAGTGGA3’ and 5’TGGTGGAACGAGGAATAACAG3’; 5’CTGGTAGCATTGAAGCTCCAG3’ and 5’TATTGGCAGCTCTGTTATTCCT3’; 5’CTATAGACAGGCTCTGGAGCT3’ and 5’AATGCCAACTTTGTATAATAAAGTTGTTTACTTGTGCGAAGAAGTTGCT3’). The resulting amplicons had matching overlaps between each other and to a PCR fragment containing an R2-L3 vector backbone (amplified with primers 5’CATAACCACTTTGTACAAGAAAGCTGA3’ and 5’AACAACTTTATTATACAAAGTTGGCAT3’) to assemble the final 4 molecules through Gibson Assembly into the entry clone.

*ProSWEET10* (5′CCTAAAGATGATAGTATTGATGA3’ forward and 5′TTTTATATCTCTCTCAAAGTAGTC3’ reverse) and *ProINV4:INV4* (5′ATTTCTCGGCTGCAAAACAT3’ forward and 5′AAGAGCTCCATCATTCATTTGCAG3’ reverse) were PCR amplified and cloned into *pBGWFS7*.*0* (*54*) and *pMDC83* (*55*) binary vectors, respectively, using the GATEWAY technology according to the manufacturer’s instructions (Thermo Fisher Scientific).

### *In vitro* culture

The solid medium for in vitro culture contained the MS basal salt mixture including vitamins (M0222, Duchefa), 0.8% (w/v) plant agar, 0.05% (w/v) MES-KOH (pH 5.8) and was supplemented with different sugars as indicated in the figures. The broad-spectrum biocide for plant tissue culture Plant Preservative Mixture (PPM, Plant Cell Technology), was supplemented to suppress airborne, waterborne, or endogenous microbial contamination, thereby minimizing the need for multiple external sterilization steps and reducing the risk of mechanical damage.

Arabidopsis flower buds at 0 days after flowering were emasculated, and one or two days after emasculation, pistils were inserted into the solid ovule-culture medium described above, supplemented with different sugars as indicated in the figures. Plates were sealed with Millipore tape and incubated under long-day conditions (16 hours light/8 hours dark) at 22°C. For *Brassica napus*, pistils were placed directly into the *in vitro* culture medium immediately after emasculation.

### *In planta* sucrose treatment

Inflorescences with emasculated flowers were immersed, 2DAE, in a sucrose or sorbitol solution and maintained in it continuously for the duration specified in the figure.

### Inducible gene expression system

*ITPS1;tps1* plants were grown in soil and sprayed with a 1 µM dexamethasone (Sigma) solution containing 0.02% (v/v) Tween-20, starting 10 days after sowing. Spraying was repeated 3-4 times at around 5 days intervals until flowering, after which induction was stopped.

*ICALS3* emasculated flowers, 0DAF, were dipped in a 100 µM estradiol (CAYMAN) or DMSO (control) solution containing 0.02% Tween-20 and, 1DAE, inserted in the solid ovule-culture medium (see above) supplemented with either 100 µM estradiol or DMSO.

*IRNAi-TOR* and *IGUS* inflorescences were dipped in a 50% (v/v) ethanol and 0.05% (v/v) Tween80 solution. In addition, an open Eppendorf tube filled with pure ethanol was placed in each pot and the plant was covered with a perforated plastic bag.

### Hormone treatment

Pistils 2DAE were dipped in a 2,4-D (Sigma) or GA3 (Sigma) 200 µM solution containing 0.05% (v/v) Tween80 as previously described (*7*). For *in vitro* culture, treated pistils were inserted in the solid ovule-culture medium (see above).

### TOR inhibitor treatment

Pistils 2DAE were inserted in the solid ovule-culture medium (see above) supplemented with 5 µM AZD8055 or 10 µM Torin2.

### Sugar measurement

Sugar quantifications in Supplementary figure 1 were performed from more than 100 ovules or seeds from two pistils or siliques per sample, as previously described (*28*). Eight biological replicates from independent plants were conducted.

Sugar quantifications in Figure 4D were performed from more than 100 ovules or seeds from two pistils or siliques per sample by LC-MS/MS, as previously described (*36*) with modifications as described by (*56*). Four biological replicates from independent plants were conducted.

Trehalose 6-phosphate was quantified from more than 50 ovules or seeds from two pistils or siliques per sample by LC-MS/MS, as previously described (*36*) with modifications as described by (*56*). Four biological replicates from independent plants were conducted.

### Lugol staining to detect starch

Harvested seeds and ovules were incubated in a 1% (w/v) SDS, 0.2 N NaOH solution at 37°C for 2 hours to clear the tissues. Samples were rinsed in water and incubated in a 12.5% bleach solution (1.25% active Cl-v/v) for 10-30 minutes. Samples were then rinsed in water and transferred to a 33% (v/v) Lugol solution in water and incubated for 30 seconds before being analyzed by differential interference contrast microscopy with an Axioplan 2 microscope (Zeiss). More than 50 ovules or seeds from three or more pistils or siliques, collected from independent plants, were analyzed per sample.

### Vanillin staining to detect proanthocyanidins

Harvested seeds and ovules were incubated in a 1% (w/v) vanillin and 6N HCl solution at room temperature for 10 minutes and analyzed by differential interference contrast microscopy with an Axioplan 2 microscope (Zeiss). More than 50 ovules or seeds from three or more pistils or siliques, collected from independent plants, were analyzed per sample.

### SR2200 staining

Harvested seeds and ovules were stained with SR2200 (Renaissance Chemicals) as previously described (*57*). More than 50 ovules or seeds from three or more pistils or siliques, collected from independent plants, were analyzed per sample.

### GUS assay

Harvested seeds and ovules were GUS stained as previously described (*9*) and analyzed by differential interference contrast microscopy with an Axioplan 2 microscope (Zeiss). More than 50 ovules or seeds from three or more pistils or siliques, collected from independent plants, were analyzed per sample.

### Tracing symplastic movement

Dissected pistils and siliques were inserted in a solid MS medium supplemented with a 0.5 mg/ml 5(6)-carboxyfluorescein diacetate solution (Invitrogen). Samples were observed after an incubation period of 3-4 hours by confocal laser scanning microscopy (Leica SP8). More than 50 ovules or seeds from three or more pistils or siliques, collected from independent plants, were analyzed per sample.

### Confocal microscopy

Fluorescent protein-expressing lines were analyzed one hour after mounting the samples in a 100 µg/mL propidium iodide and 7% (w/v) sucrose solution, as previously described (*7*). Samples were imaged by confocal laser scanning microscopy (Leica SP8), and analyzed with the ImageJ software (*58*).

More than 50 ovules or seeds from three or more pistils or siliques, collected from independent plants, were analyzed per sample.

The mean fluorescence intensity of five cells per sample was quantified in at least seven ovules or seeds from three or more pistils or siliques, collected from independent plants, using the ImageJ software.

Corrected total cell fluorescence was calculated as Integrated Density – (Area of selected cells x Mean fluorescence of background readings) in at least 14 ovules or seeds from three pistils or siliques, collected from independent plants, using the ImageJ software.

### Transcriptomic analyses

Libraries were prepared for Columbia-0 siliques at 0 and 2 DAF (three replicates for each genotype and time point) according to the Illumina TruSeq RNA protocol and sequenced in two lanes of a Novaseq 6000 system, yielding an average of 70M read pairs per sample. Reads for each biological replicate were aligned independently to the Arabidopsis thaliana TAIR10 reference genome using hisat2 v2.1.0 (*59*), allowing a maximum intron length of 57700bp (the largest intron in the TAIR10 annotation). An average of 81.9% of the reads aligned to the reference (minimum of 77.7%). The number of reads per transcript in the TAIR10 annotation was counted with the featureCounts function in the Rsubread R package with default parameters (*60*). Only transcripts that presented more than 10 reads in all samples together were used in downstream analysis, leaving us with 25053 transcripts out of the 32678 transcripts present in the annotation.

Sample homogeneity was surveyed in R with the plotPCA function in the DEseq2 package (*61*) and with the PoissonDistance function in the Bioconductor PoiClaClu package (*62*).

Gene ontology (GO) enrichment analyses were conducted on the ShinyGO website with an FDR cutoff < 0.05 (*63*).

Heatmap analyses were conducted on the Morpheus website (https://software.broadinstitute.org/morpheus). Hierarchical clustering was performed with the One Minus Pearson’s Correlation metric and average linkage method.

### Accession Numbers

AGL62 (AT5G60440), FIE (AT3G20740), TPS1 (AT1G78580), TOR (AT1G50030), CALS6 (AT3g59100), CALS7 (AT1G06490), SWEET10 (AT5G50790), INV4 (AT2G36190), RGA (AT2G01570), GAI (AT1G14920).

## Discussion

Vivipary, the process by which embryos develop within the parent, had a profound impact on the evolution of life on our planet. Whereas the phenomenon pertains to both animal and plants, the term has been used in plants solely to describe seeds that germinate while still attached to the plant. Nevertheless, seeds have a maternal component - the seed coat, the funiculus, the chalaza, and the nucellus - that protects and nourishes the embryo regardless of germination. This evolutionary advantage was made possible by a signaling pathway for zygote-maternal communication. The animal embryo and the plant endosperm initiate signals that, despite their different evolutionary origin and chemical nature, trigger a fundamentally similar maternal response in charge of accommodating and nourishing the growth of the embryo. Nonetheless, we highlighted here how plants integrate developmental signals with their energy status. We showed that maternal sucrose, necessary as source of carbon and energy for the growth of the seed, carries the signal information responsible for the development of maternal tissues. By contrast, it did not stimulate the growth of the central and egg cell, thus implying the presence of a zygotic checkpoint in the development of the seed. An increase in the content of soluble sugars, among which sucrose, was induced by fertilization in an AGL62 dependent manner and correlated with a change in the expression level of sucrose and starch metabolic genes. We demonstrated that fertilization transform the seed into a sugar-sink by facilitating both the symplastic and apoplastic transport of sucrose. Before fertilization, callose synthases CALS 6 and 7 are expressed in the funiculus and chalaza to restrict the influx of sugars into the ovule. Fertilization activates glucanases with callose hydrolase activity, enhancing the flow of sugars into developing seeds (*27*). Our data show that the symplastic movement into the seed is facilitated by T6P, a signaling molecule synthetized in response to fertilization. Mutations in the *TPS1* gene, encoding the main enzyme responsible for T6P synthesis in the funiculus and chalaza, inhibited maternal response and starch accumulation in the seed, similar to *agl62* seeds. T6P production was also stimulated by high sucrose levels, thus creating a positive feed-back loop that likely sustains further sucrose transfer into the seed. Fertilization also induced the expression of a key module for sucrose apoplastic transport, comprising the sucrose transporter SWEET10, which facilitates sucrose export to the apoplast, and the cell wall invertase INV4, which hydrolyzes sucrose into hexoses. Both genes were expressed in the chalaza and activated by auxin, which, as predicted, led to increased levels of sucrose, glucose and fructose. Auxin stimulated also the synthesis of T6P, but only in the presence of sucrose. We propose that auxin drives T6P production by elevating sucrose levels through enhanced apoplastic transport, thus further facilitating sugar flow. Both auxin and T6P signaling pathways are repressed by the PcG protein FIE, which prevents a fertilization-independent response of maternal tissues. Our results suggest that fertilization releases the FIE repressive mechanism, thus activating auxin and T6P signaling to facilitate both apoplastic and symplastic sucrose transport into the seed. The sugar signal is than transduced into a gibberellin response that stimulates the biosynthesis of PAs. The application of sugar led to the elimination of the RGA DELLA protein, a negative regulator of the gibberellin signaling pathway, while overexpression of the GAI DELLA protein blocked the synthesis of PAs without affecting starch accumulation. The gibberellin pathway required the presence of TOR in order to stimulate PA deposition. The inhibition of TOR function or expression did not affect starch accumulation but blocked the maternal response to sucrose regardless of the presence of GA, thus highlighting its central role in the fertilization pathway. It has been shown that TOR regulates FIE function in the vernalization-induced floral transition (*64*), implying a potentially similar interaction in the seed.

## Acknowledgements

We thank Yrjo Helariutta for providing the ICALS3 line and the Observatoire du Végétal for plant culture, access to imaging and mass spectrometry facilities and assistance.

## Authors contributions

W.X. designed and performed the research and analyzed the data. M.I. performed genetic and expression analyses and hormone treatment experiments. D.M.G.P. and J.L. helped to characterize SWEET10 and INV4. S.C. performed genetic analyses and assisted al research aspects. R.L.H. ad S.D. performed sugar quantification analyses and helped to design, analyze, and interpret sugar transport results. L.K. created the *CALS6* promoter-reporter and *cals6;cals7* double mutant line. R.F. and J.E.L. quantified T6P and helped to design, analyze, and interpret T6P results. C.M. helped to design, analyze, and interpret TOR results. E.M. designed the research, analyzed the data and wrote the article.

## References

1. H. B. Braz et al., Evolutionary Patterns of Maternal Recognition of Pregnancy and Implantation in Eutherian Mammals. Animals (Basel) 14, (2024).

2. P. Duc-Goiran, T. M. Mignot, C. Bourgeois, F. Ferré, Embryo-maternal interactions at the implantation site: a delicate equilibrium. Eur J Obstet Gynecol Reprod Biol 83, 85–100 (1999).

3. P. C. K. Leung, E. Y. Adashi, The Ovary. (Academic Press, ed. Third Edition, 2019).

4. R. J. Camerarius. (1694).

5. N. Grew, The anatomy of plants : with an idea of a philosophical history of plants; and several other lectures read before the Royal Society. (Printed by W. Rawlins for the author, London, 1682).

6. G. N. Drews, R. Yadegari, Development and function of the angiosperm female gametophyte. Annu Rev Genet 36, 99–124 (2002).

7. D. D. Figueiredo, R. A. Batista, P. J. Roszak, L. Hennig, C. Köhler, Auxin production in the endosperm drives seed coat development in. Elife 5, (2016).

8. P. Roszak, C. Köhler, Polycomb group proteins are required to couple seed coat initiation to fertilization. Proc Natl Acad Sci U S A 108, 20826–20831 (2011).

9. W. Xu et al., Endosperm and Nucellus Develop Antagonistically in Arabidopsis Seeds. Plant Cell 28, 1343–1360 (2016).

10. J. F. Gutierrez-Marcos, M. Constância, G. J. Burton, Maternal to offspring resource allocation in plants and mammals. Placenta 33 Suppl 2, e3–10 (2012).

11. F. L. Lopes et al., Sugar signaling modulates SHOOT MERISTEMLESS expression and meristem function in. Proc Natl Acad Sci U S A 121, e2408699121 (2024).

12. L. S. Meng et al., Glucose- and sucrose-signaling modules regulate the Arabidopsis juvenile-to-adult phase transition. Cell Rep 36, 109348 (2021).

13. S. Smeekens, SUGAR-INDUCED SIGNAL TRANSDUCTION IN PLANTS. Annu Rev Plant Physiol Plant Mol Biol 51, 49–81 (2000).

14. H. Weber, L. Borisjuk, U. Wobus, Molecular physiology of legume seed development. Annu Rev Plant Biol 56, 253–279 (2005).

15. T. J. Ross-Elliott et al., Phloem unloading in Arabidopsis roots is convective and regulated by the phloem-pole pericycle. Elife 6, (2017).

16. B. Riens, G. Lohaus, D. Heineke, H. W. Heldt, Amino Acid and sucrose content determined in the cytosolic, chloroplastic, and vacuolar compartments and in the Phloem sap of spinach leaves. Plant Physiol 97, 227–233 (1991).

17. R. Stadler, C. Lauterbach, N. Sauer, Cell-to-cell movement of green fluorescent protein reveals post-phloem transport in the outer integument and identifies symplastic domains in Arabidopsis seeds and embryos. Plant Physiol 139, 701–712 (2005).

18. L. Q. Chen et al., A cascade of sequentially expressed sucrose transporters in the seed coat and endosperm provides nutrition for the Arabidopsis embryo. Plant Cell 27, 607–619 (2015).

19. N. Grignon, B. Touraine, M. Durand, 6(5)Carboxyfluorescein as a Tracer of Phloem Sap Translocation. American Journal of Botany 76, 871–877 (1989).

20. C. Faulkner, Plasmodesmata and the symplast. Curr Biol 28, R1374–R1378 (2018).

21. M. F. Belmonte et al., Comprehensive developmental profiles of gene activity in regions and subregions of the Arabidopsis seed. Proc Natl Acad Sci U S A 110, E435–444 (2013).

22. D. H. Barratt et al., Callose synthase GSL7 is necessary for normal phloem transport and inflorescence growth in Arabidopsis. Plant Physiol 155, 328–341 (2011).

23. B. Xie, X. Wang, M. Zhu, Z. Zhang, Z. Hong, CalS7 encodes a callose synthase responsible for callose deposition in the phloem. Plant J 65, 1–14 (2011).

24. L. Záveská Drábková, D. Honys, Evolutionary history of callose synthases in terrestrial plants with emphasis on proteins involved in male gametophyte development. PLoS One 12, e0187331 (2017).

25. L. Kalmbach et al., Putative pectate lyase PLL12 and callose deposition through polar CALS7 are necessary for long-distance phloem transport in Arabidopsis. Curr Biol 33, 926–939.e929 (2023).

26. A. Vatén et al., Callose biosynthesis regulates symplastic trafficking during root development. Dev Cell 21, 1144–1155 (2011).

27. X. Liu et al., Fertilization-dependent phloem end gate regulates seed size. Curr Biol 35, 2049–2063.e2043 (2025).

28. J. Lu et al., The nucellus: between cell elimination and sugar transport. Plant Physiol 185, 478–490 (2021).

29. L. Q. Chen et al., Sucrose efflux mediated by SWEET proteins as a key step for phloem transport. Science 335, 207–211 (2012).

30. T. Roitsch, M. C. González, Function and regulation of plant invertases: sweet sensations. Trends Plant Sci 9, 606–613 (2004).

31. J. W. Riesmeier, L. Willmitzer, W. B. Frommer, Isolation and characterization of a sucrose carrier cDNA from spinach by functional expression in yeast. EMBO J 11, 4705–4713 (1992).

32. N. Sauer, J. Stolz, SUC1 and SUC2: two sucrose transporters from Arabidopsis thaliana; expression and characterization in baker’s yeast and identification of the histidine-tagged protein. Plant J 6, 67–77 (1994).

33. J. R. Gottwald, P. J. Krysan, J. C. Young, R. F. Evert, M. R. Sussman, Genetic evidence for the in planta role of phloem-specific plasma membrane sucrose transporters. Proc Natl Acad Sci U S A 97, 13979–13984 (2000).

34. F. Andrés et al., The sugar transporter SWEET10 acts downstream of FLOWERING LOCUS T during floral transition of Arabidopsis thaliana. BMC Plant Biol 20, 53 (2020).

35. J. M. Ruhlmann, B. W. Kram, C. J. Carter, CELL WALL INVERTASE 4 is required for nectar production in Arabidopsis. J Exp Bot 61, 395–404 (2010).

36. J. E. Lunn et al., Sugar-induced increases in trehalose 6-phosphate are correlated with redox activation of ADPglucose pyrophosphorylase and higher rates of starch synthesis in Arabidopsis thaliana. Biochem J 397, 139–148 (2006).

37. A. J. van Dijken, H. Schluepmann, S. C. Smeekens, Arabidopsis trehalose-6-phosphate synthase 1 is essential for normal vegetative growth and transition to flowering. Plant Physiol 135, 969–977 (2004).

38. L. D. Gómez, A. Gilday, R. Feil, J. E. Lunn, I. A. Graham, AtTPS1-mediated trehalose 6-phosphate synthesis is essential for embryogenic and vegetative growth and responsiveness to ABA in germinating seeds and stomatal guard cells. Plant J 64, 1–13 (2010).

39. F. Fichtner, J. E. Lunn, The Role of Trehalose 6-Phosphate (Tre6P) in Plant Metabolism and Development. Annu Rev Plant Biol 72, 737–760 (2021).

40. F. Fichtner et al., Functional Features of TREHALOSE-6-PHOSPHATE SYNTHASE1, an Essential Enzyme in Arabidopsis. Plant Cell 32, 1949–1972 (2020).

41. C. Y. Liao et al., Reporters for sensitive and quantitative measurement of auxin response. Nat Methods 12, 207–210, 202 p following 210 (2015).

42. D. D. Figueiredo, R. A. Batista, P. J. Roszak, C. Köhler, Auxin production couples endosperm development to fertilization. Nat Plants 1, 15184 (2015).

43. C. Köhler, G. Makarevich, Epigenetic mechanisms governing seed development in plants. EMBO Rep 7, 1223–1227 (2006).

44. M. Oliva et al., FIE, a nuclear PRC2 protein, forms cytoplasmic complexes in Arabidopsis thaliana. J Exp Bot 67, 6111–6123 (2016).

45. E. Dorcey, C. Urbez, M. A. Blázquez, J. Carbonell, M. A. Perez-Amador, Fertilization-dependent auxin response in ovules triggers fruit development through the modulation of gibberellin metabolism in Arabidopsis. Plant J 58, 318–332 (2009).

46. S. Feng et al., Coordinated regulation of Arabidopsis thaliana development by light and gibberellins. Nature 451, 475–479 (2008).

47. B. Menand et al., Expression and disruption of the Arabidopsis TOR (target of rapamycin) gene. Proc Natl Acad Sci U S A 99, 6422–6427 (2002).

48. Y. Xiong et al., Glucose-TOR signalling reprograms the transcriptome and activates meristems. Nature 496, 181–186 (2013).

49. D. Deprost et al., The Arabidopsis TOR kinase links plant growth, yield, stress resistance and mRNA translation. EMBO Rep 8, 864–870 (2007).

50. C. M. Chresta et al., AZD8055 is a potent, selective, and orally bioavailable ATP-competitive mammalian target of rapamycin kinase inhibitor with in vitro and in vivo antitumor activity. Cancer Res 70, 288–298 (2010).

51. Q. Liu et al., Discovery of 9-(6-aminopyridin-3-yl)-1-(3-(trifluoromethyl)phenyl)benzo[h][1,6]naphthyridin-2(1H)-one (Torin2) as a potent, selective, and orally available mammalian target of rapamycin (mTOR) inhibitor for treatment of cancer. J Med Chem 54, 1473–1480 (2011).

52. M. H. Montané, B. Menand, ATP-competitive mTOR kinase inhibitors delay plant growth by triggering early differentiation of meristematic cells but no developmental patterning change. J Exp Bot 64, 4361–4374 (2013).

53. M. A. Salem, Y. Li, A. Wiszniewski, P. Giavalisco, Regulatory-associated protein of TOR (RAPTOR) alters the hormonal and metabolic composition of Arabidopsis seeds, controlling seed morphology, viability and germination potential. Plant J 92, 525–545 (2017).

54. M. Karimi, D. Inzé, A. Depicker, GATEWAY vectors for Agrobacterium-mediated plant transformation. Trends Plant Sci 7, 193–195 (2002).

55. M. D. Curtis, U. Grossniklaus, A gateway cloning vector set for high-throughput functional analysis of genes in planta. Plant Physiol 133, 462–469 (2003).

56. C. M. Figueroa et al., Trehalose 6-phosphate coordinates organic and amino acid metabolism with carbon availability. Plant J 85, 410–423 (2016).

57. D.-M. Gómez-Páez, E. Magnani, in Seed Dormancy: Methods and Protocols, N. Kawakami, K. Sato, Eds. (Springer US, New York, NY, 2024), pp. 93–104.

58. C. A. Schneider, W. S. Rasband, K. W. Eliceiri, NIH Image to ImageJ: 25 years of image analysis. Nat Methods 9, 671–675 (2012).

59. D. Kim, B. Langmead, S. L. Salzberg, HISAT: a fast spliced aligner with low memory requirements. Nat Methods 12, 357–360 (2015).

60. Y. Liao, G. K. Smyth, W. Shi, The R package Rsubread is easier, faster, cheaper and better for alignment and quantification of RNA sequencing reads. Nucleic Acids Res 47, e47 (2019).

61. M. I. Love, W. Huber, S. Anders, Moderated estimation of fold change and dispersion for RNA-seq data with DESeq2. Genome Biol 15, 550 (2014).

62. W. Dm, Classification and clustering of sequencing data using a Poisson model. Annals of Applied Statistics 5, 2493–2518 (2011).

63. S. X. Ge, D. Jung, R. Yao, ShinyGO: a graphical gene-set enrichment tool for animals and plants. Bioinformatics 36, 2628–2629 (2020).

64. R. Ye et al., Glucose-driven TOR-FIE-PRC2 signalling controls plant development. Nature 609, 986–993 (2022).

