## Supplemental Figures for "Too much sugar makes plants ‘pregnant’: maternal sucrose signals fertilization in Arabidopsis seeds"

**Supplementary figures**

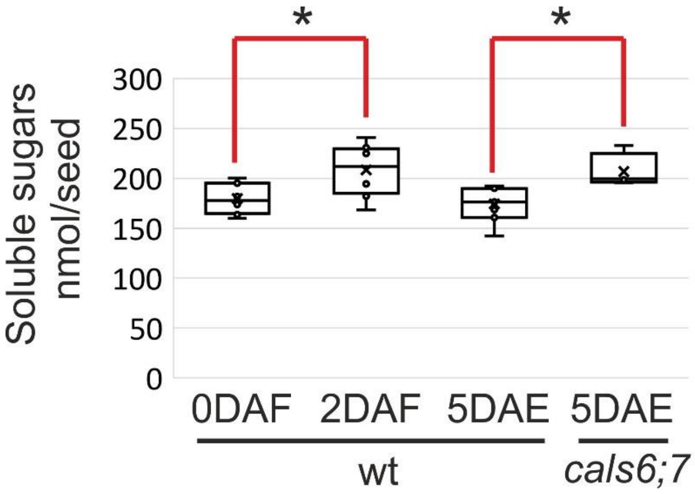


**Supplementary figure 1**

Quantification of soluble sugars (sucrose, glucose, and fructose) levels in wild type and *cals6;7* ovules and seeds. In the box and whisker chart, the rectangles represent the interquartile ranges (median excluded), the mid-lines represent the medians and the whiskers represent the minimum and maximum values within 1.5 times the interquartile range. Asterisks indicate statistically significant difference (Student's t-test, P<0.05, n=8 biological replicates).

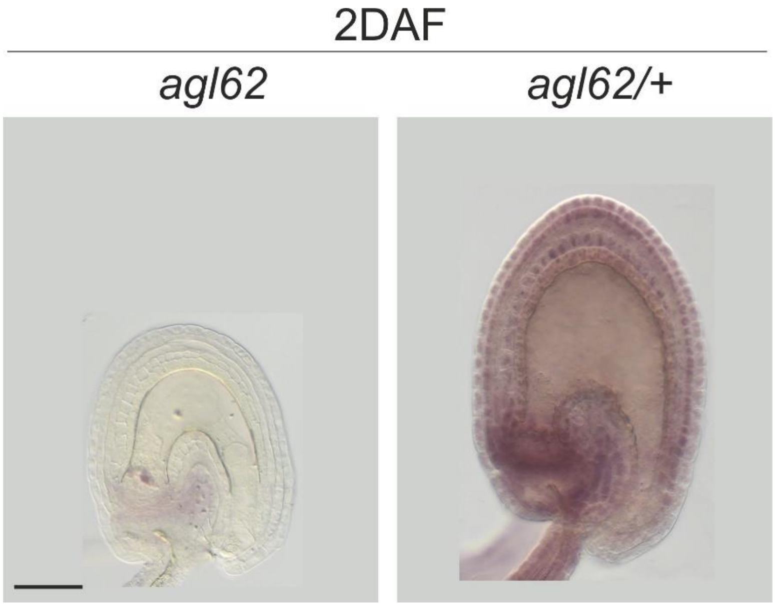


**Supplementary Figure 2.**

Differential interference contrast microscopy images of seeds carrying an agl62 or agl62/+ endosperm stained with Lugol’s solution to show the presence of starch. DAF, days after flowering. Bar = 50 µm.

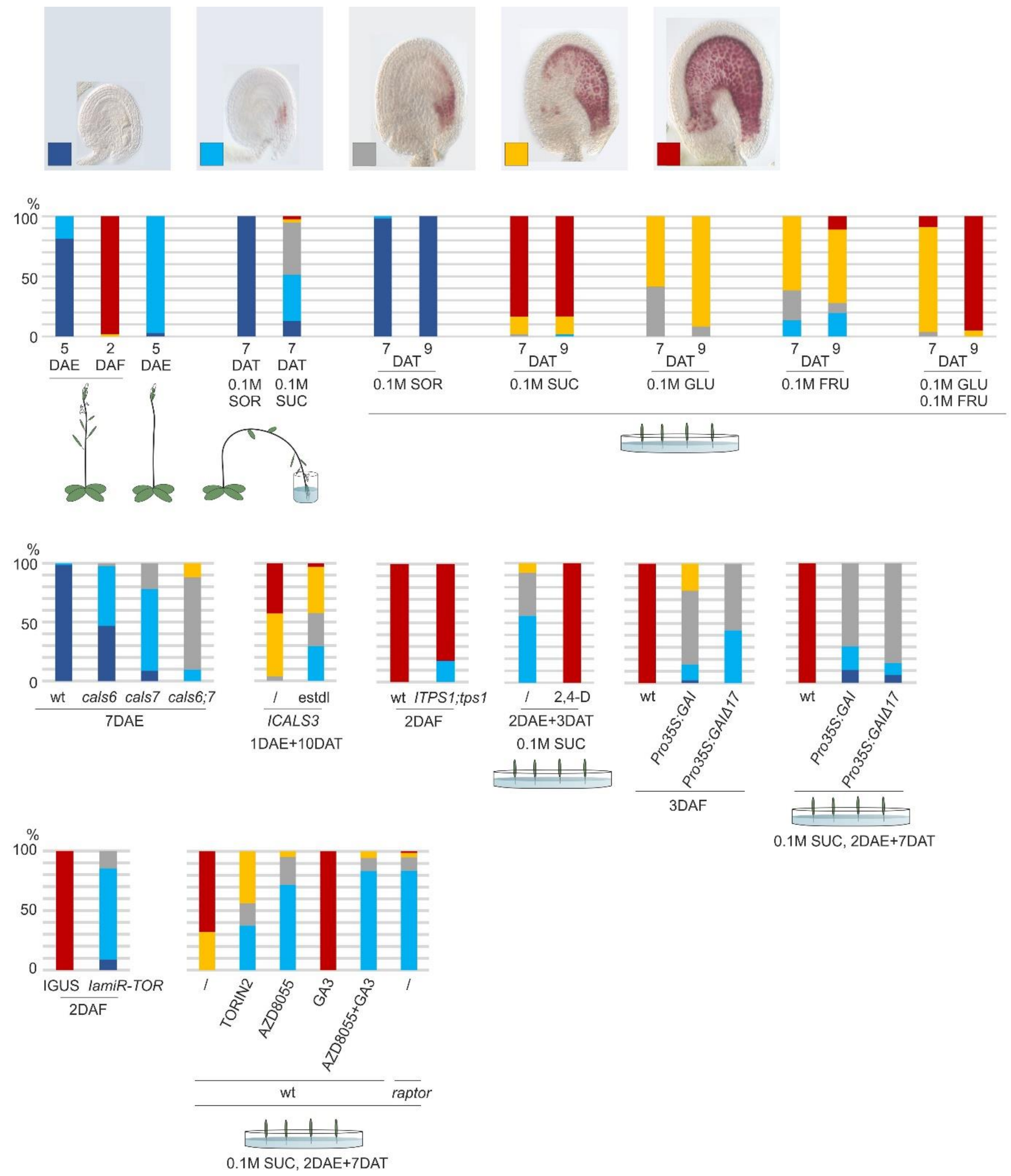


**Supplementary Figure 3.**

Visual quantification of vanillin staining, expressed as the percentage of seeds exhibiting a staining pattern consistent with the legend (blue: no staining; cyan: staining in the micropyle; grey: staining till the bending zone; yellow: staining beyond the bending zone; red: full staining). More than 50 ovules or seeds from three or more pistils or siliques, collected from independent plants, were analyzed per sample. /, mock treatment; wt; wild type; DAE, days after emasculation; DAF, days after flowering; DAT, days after treatment; SUC, sucrose; SOR, sorbitol; GLU, glucose; FRU, fructose.

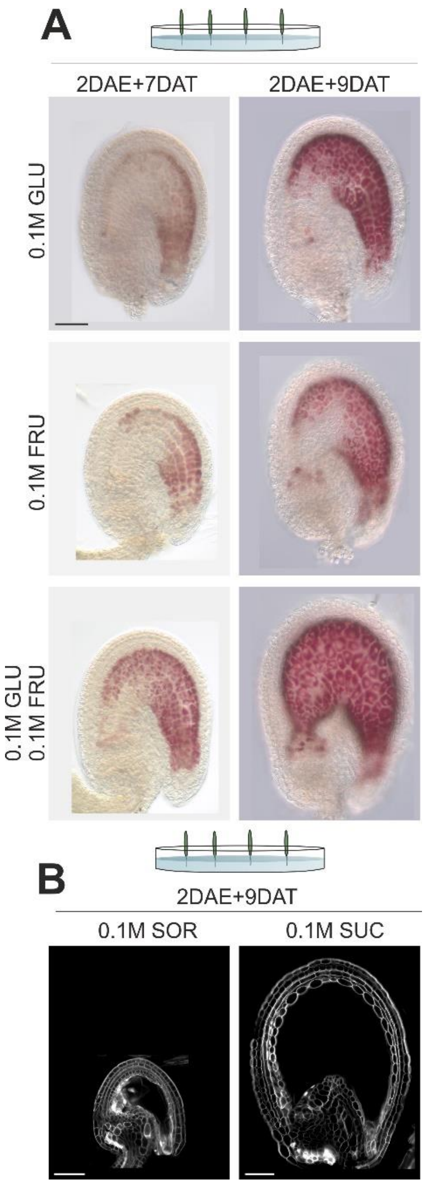


**Supplementary Figure 4.**

**(A)** DIC images of ovules stained with vanillin. **(B)** Confocal images of ovules stained with SR2200. DAE, days after emasculation; DAT, days after treatment; GLU, glucose; FRU, fructose; SOR, sorbitol; SUC, sucrose; Bars = 50 µm.

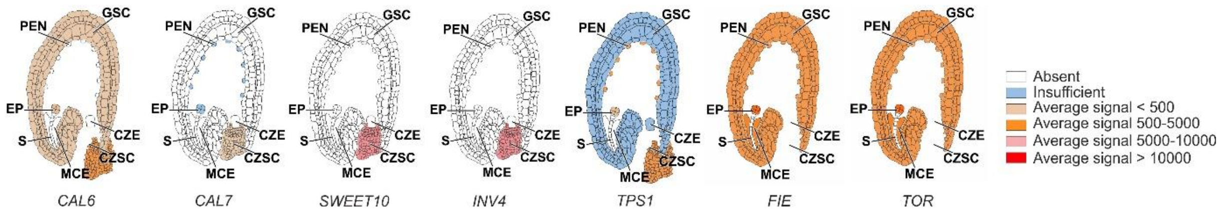


**Supplementary Figure 5.**

GeneChip Expression Profile of genes in different domains of the seed at the pre-globular embryo stage as obtained by laser capture microdissection (*21*). The colors correspond to the average signal intensity of biological replicates (from white, no signal, to red, signal greater than 10000). CZE, chalazal endosperm; CZSC, chalazal seed coat; EP, embryo proper; GSC, general seed coat; MCE, micropylar endosperm; PEN, peripheral endosperm; S, suspensor.

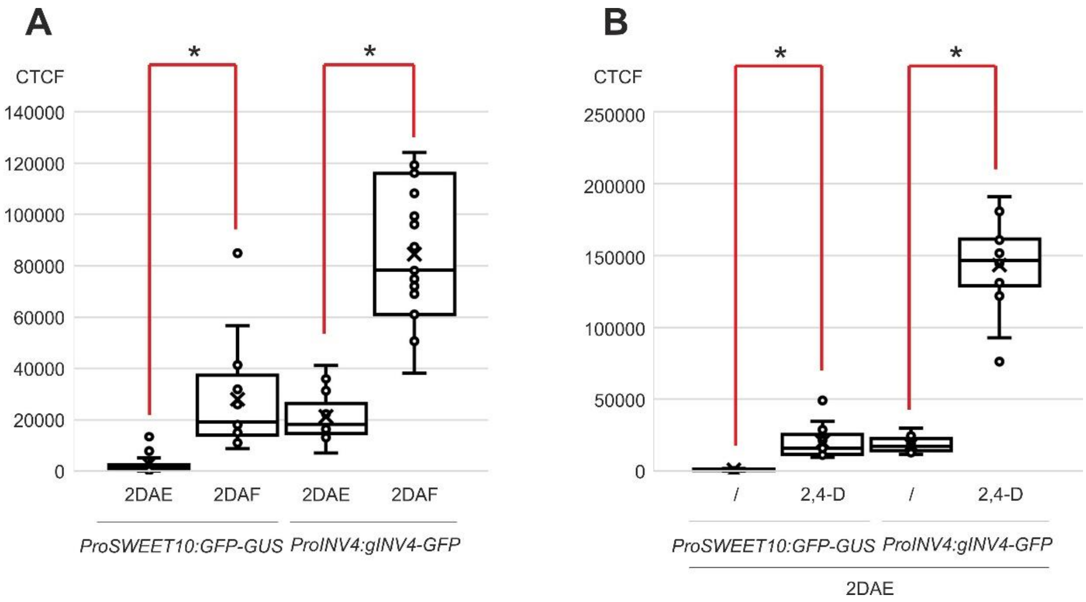


**Supplementary Figure 6.**

**(A)** Quantification of GFP signal in *proSWEET10:GFP-GUS* and *proINV4:gINV4-GFP* unfertilized ovules and seeds. **(B)** Quantification of GFP signal in *proSWEET10:GFP-GUS* and *proINV4:gINV4-GFP* unfertilized ovules treated with the synthetic auxin 2,4-D. In the box and whisker chart, the rectangles represent the interquartile ranges (median excluded), the mid-lines represent the medians and the whiskers represent the minimum and maximum values within 1.5 times the interquartile range. Asterisks indicate statistically significant difference (Student's t-test, $P<0.05$, $n>14$). DAE, days after emasculation; DAF, days after flowering. CTCF, corrected total cell fluorescence.
